# Manual gestures facilitate learning lexical stress by modulating auditory neural responses

**DOI:** 10.1101/2023.11.14.566652

**Authors:** Tianqi Zhan, Danfeng Yang, Ruoyu Wu, Xing Tian

## Abstract

Gestures accompany speech and facilitate communication and learning. Most previous studies have demonstrated the effects of gestures on learning semantics, yet how gestures facilitate learning low- and intermediate-level speech features is unclear. The present study investigated the effects of manual gestures on learning lexical stress, a phonological-lexical feature that is foreign to native Mandarin speakers. Across a series of experiments, we demonstrated that the gestures with representational relations to auditory stimuli in terms of covaried amplitude modulation facilitated the learning of lexical stress for both familiar (English) and unknown (Russian) languages, but not for pseudowords that lack phonotactic properties. Interestingly, gestures with amplitude trajectory matching the stress benefited the learning of trained words, whereas gestures that only matched the timing of syllable segments but not amplitude variation generalized the learning effects to untrained stimuli. Furthermore, in the EEG experiment, we found that gesture-accompanied learning was associated with power increase and inter-trial phase coherence (ITC) decrease in the theta band at the time windows corresponding to the stress positions. These results suggest that the facilitatory effects of gestures on lexical stress learning depend on the specificity of cross-modal feature mapping at the phonological level, mediated by the neural modulation in early perceptual responses.

## 1 Introduction

When engaging in conversations, our hands often move naturally in company with our speech, either spontaneously or intentionally. Such co-speech gestures have been found to significantly aid in the production and comprehension of language (Driskell & Radtke, 2003; Macedonia, 2014; McNeill, 2007).

With the increasing popularity of embodied learning during the past few decades (Mathias & Kriegstein, 2023), the advantage of this well-established link between language and gestures has been extensively demonstrated in language acquisition. A robust facilitation effect of gestures in semantic learning has been observed across language units, paradigms, and demographic groups (Argyriou et al., 2017; Dargue et al., 2019; Goldin-Meadow & Wagner, 2005; Hadar et al., 1998; Macedonia & Knösche, 2011)

However, compared to the studies in semantic learning, fewer studies investigate the role of gestures in learning low- and intermediate-level speech features that are crucial in linking surface structures and semantics, and the observations from these studies have been inconsistent. Some studies have provided support for the facilitation effects of gestures in learning some speech attributes. For example, evidence in several studies demonstrate that directional gestures help non-native Chinese speakers in learning Mandarin tones (Baills et al., 2019; Morett & Chang, 2015; Zhen et al., 2019). In contrast, Jesse and Mitterer (2011) did not find any impacts of pointing gestures on the perception of Dutch lexical stress. Hirata and Kelly (2010) found that gestures did not enhance the ability of native English speakers to discriminate the length of Japanese vowels.

One possibility to account for these inconsistent findings is that the learning effect depends on the nature of the relations between gestures and speech stimuli. For example, the relationship between these two sources of stimuli needs to be consistent with one’s prior experience or natural settings in some *general* aspects, such as concordance of occurrence in space and time (Shams & Seitz, 2008). Moreover, congruency is also important at a more *specific* featural level, evident by studies showing that multisensory training can induce more pronounced effects using featural congruent stimuli than using arbitrary pairing of stimuli (Bavelier & Neville, 2002; Kim et al., 2008; Macedonia & Knösche, 2011). These results suggest that the mapping between the features of gestures and acoustic stimuli may provide a bridge across modalities and facilitate learning by gestures. The *generality vs. specificity* in the relations across modalities may constrain the facilitation effects of gestures on speech learning.

In the present study, we carried out a series of behavioral and EEG experiments to investigate how gestures affect the learning of intermediate-level speech features using lexical stress as a research model. Lexical stress refers to the emphasis or prominence placed on certain syllables within a word. In English, lexical stress is characterized by a greater amplitude, higher fundamental frequency (F0), and longer syllable duration (Lieberman, 1960), functioning as an important cue in spoken word recognition (Jesse et al., 2017). However, lexical stress is a distinctive lexical-phonological feature of English but not Mandarin, which poses challenges for Mandarin speakers in perceiving and learning English lexical stress compared to other English learners (Archibald, 1997). Therefore, our first aim was to establish the ground truth of whether gestures can help native Mandarin speakers learn English lexical stress. We designed a learning paradigm in Behavioral Experiment 1 (BE1), where participants watched the videos of continuous up-and-down hand movements synchronized with the audio recordings of trisyllable words, with each syllable temporally aligned with one cycle of up-and-down movement. The learning effects of lexical stress were quantified by comparing the accuracies of stress position judgments before and after learning. Additionally, building upon previous findings that gestures can enhance word recognition in both the short and long terms (Macedonia & Klimesch, 2014; Masumoto et al., 2006), we also examined whether any observed immediate learning effects can persist at least a day after the initial learning session.

The second aim is to specify the functions of congruency in gesture-accompanied learning. In natural contexts, co-speech gestures typically exhibit maximum displacement coinciding with the stressed syllable (Biau & Soto-Faraco, 2013; Krahmer & Swerts, 2007) in the dimension of amplitude – a larger gesture in a repertoire shares commonality with a stressed syllable in a word. We compared the effects of four different types of gestures: *Match* (congruent), *Mismatch* (incongruent), *Balanced*, and *Static*. The congruency of the gestures is defined by whether the amplitude variation of the gesture trajectory aligns with that of the word audio. Based on the importance of congruency revealed by previous findings, we hypothesized that the *Match* gestures would facilitate the learning of lexical stress, whereas *Mismatch* gestures would not. As control groups, we also examined two other types of gestures: *Balanced* gesture, where the up-and-down amplitude of the movement identical regardless of the stress position, and *Static* gesture, where the hand remains in a fixed position. In line with the predictions from the multisensory learning framework (Shams & Seitz, 2008), we did not expect strong learning effects in these two groups given that these gestures do not provide information about the stress position.

The third aim is to test whether the learning effects can be generalized. Generalization (or transfer) of learning refers to the ability to apply knowledge or skills learned in one context to another. Embodied learning studies have demonstrated that engaging in relevant actions produces a larger generalization effect (Chang et al., 2014; Macedonia, 2019). Given the extensive functional connections between motor and sensory domains such as that movement observation can also activate motor areas (Mayer et al., 2015), it is possible that learning by simply observing co-speech gestures could also be generalized. To test this hypothesis, we assessed the performance of the stress position judgments for the unstudied words in BE1. Surprisingly, the *Balanced* gestures, but not the *Match* gestures, induced generalization one day after the initial learning session. However, since the unstudied words had already been presented to the participants during the immediate posttest, it was unclear whether the observed delayed improvement was truly due to generalization or simply a result of repeated exposure to the test stimuli. We therefore conducted Behavioral Experiment 2 (BE2) to replicate BE1 and further investigated whether *Balanced* gestures indeed induce generalization, by testing the learning effects on an additional group of new words on the second day.

Fourth, we devised Behavioral Experiments 3 (BE3) and 4 (BE4) to investigate at which level gestures help the learning of lexical stress. Specifically, we aimed to probe whether the facilitation effects of gestures depend on higher-level lexical-semantic knowledge or it may be a general increase of auditory sensitivity to lower-level acoustic attributes. In BE3, we removed any lexical-semantic information by substituting the English words with Russian words, a language that participants had little knowledge about but still possesses stress patterns characterized by enhanced amplitude at stressed syllables (Lavitskaya & Kabak, 2014). If gestures help learning by interacting with phonological processing without requiring lexical-semantic knowledge, we would still expect to observe improved test performance after learning. In BE4, we further eliminated phonotactic regularities by using pseudowords that were created by concatenating three consonant-vowel syllables. If the effects observed in BE4 are different from those in other experiments, it will suggest that the facilitation of learning happens on the abstract phonological pattern of lexical stress rather than the boost of sensitivity to lower-level acoustic attributes.

Finally, although the behavioral benefits of gestures in language learning have been extensively demonstrated, their underlying neural mechanisms are still unclear. Therefore, we conducted an electroencephalography (EEG) experiment to examine the neural correlates of gesture-accompanied learning. Unisensory perceptual learning studies have reported training-associated changes in the amplitude of early perceptual response components, such as N1/P2 complex (Carcagno & Plack, 2011; Tong et al., 2009; Wassenhove & Nagarajan, 2007). If gesture-accompanied learning modulates the phonological representation of lexical stress at the perceptual stage, we expect to see altered early auditory responses after learning. More importantly, the changes in neural responses to the words should occur at corresponding latencies of different stress positions.

## 2 Methods

### 2.1 Participants

A total of four behavioral experiments (BE) and one EEG experiment (EE) were conducted with separate groups of participants. Specifically, 124 participants participated in BE1 (105 females; mean age = 21.26 ± 2.19 years, age range = 18-28 years); 32 participants in BE2 (25 females; mean age = 21.63 ± 3.63 years, age range = 18-25 years); 31 participants in BE3 (25 females; mean age = 21.58 ± 3.58 years, age range = 18-24 years); 31 participants in BE4 (21 females; mean age = 21.88 ± 5.13 years, age range = 18-27 years); 21 participants in EE (18 females; mean age = 21.32 ± 1.83 years, age range = 18-25 years). All participants were native Mandarin speakers who had above-moderate English proficiency, with normal or corrected-to-normal vision and normal hearing. No participant reported a history of neurologic deficits. All participants signed informed consent and received monetary incentives. All protocols were approved by the Institutional Review Board at New York University Shanghai.

### 2.2 Stimuli and procedures

#### 2.2.1 Behavioral Experiment 1

##### Stimuli

Stimuli and procedures are summarized in Fig. 1. In BE1, 84 trisyllabic English words were selected from the Corpus of Contemporary American English (COCA) word frequency table, with equal numbers of words with stress on the first, second, or third syllable (28 each). Word frequencies did not significantly differ among different stress positions (frequency rank for words that stress on the 1^st^ syllable: *M* = 11835.50, *SD* = 3980.43, *range* = 8436-20199; on the 2^nd^ syllable: *M* = 11923.47, *SD* = 3906.62, *range* = 10001-20165; on the 3^rd^ syllable: *M* = 10596.25, *SD* = 2777.52, *range* = 8160-19921).

**Figure 1.**
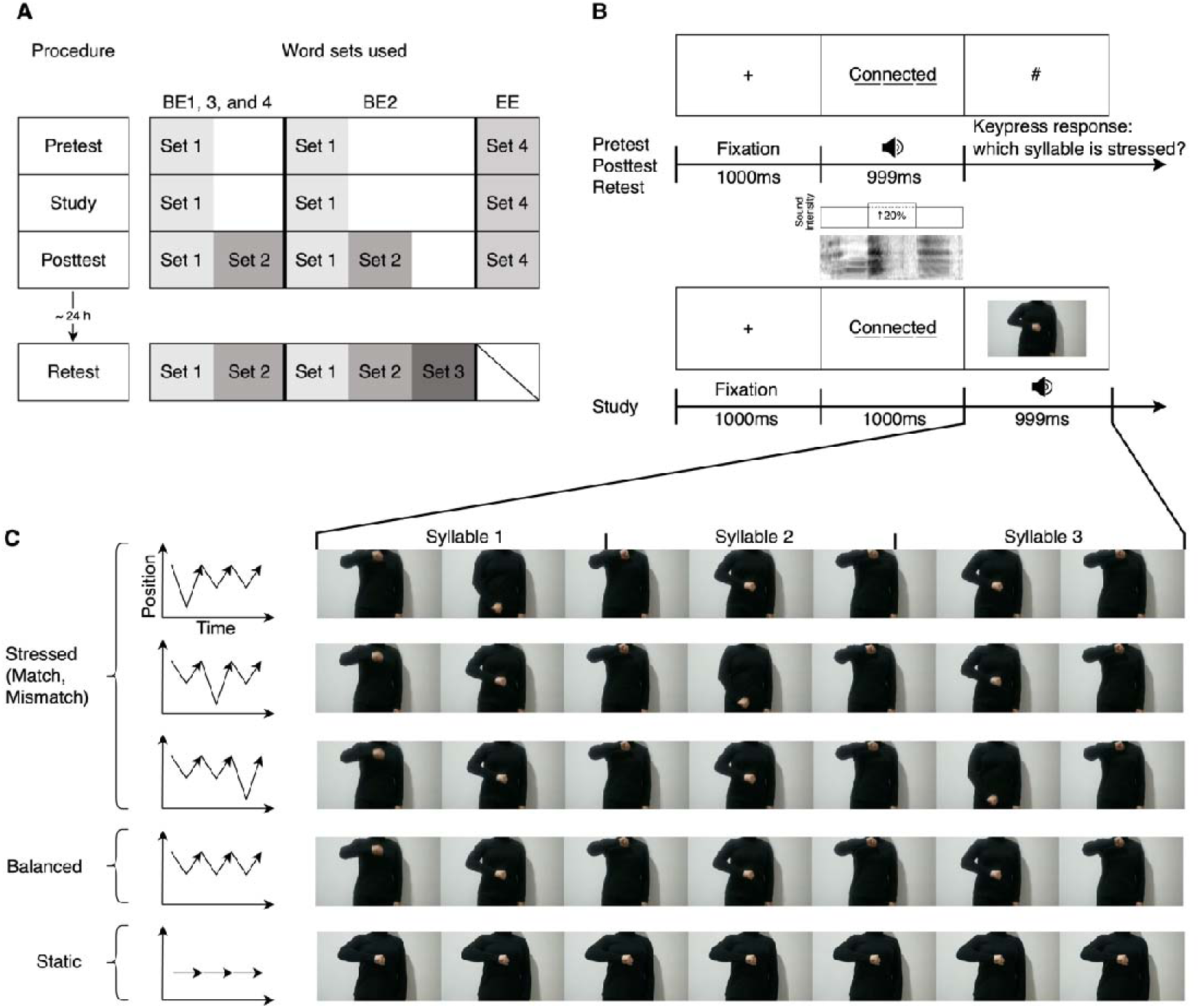
Experimental procedure and stimuli. **(A)** Summary of the experimental procedures for all experiments. The leftmost column shows the general procedure. The experiment consisted of Pretest, Study, and Posttest phases in sequence, followed by a Retest about 24 hours later. The right columns illustrate the word sets used for each phase in each experiment. Each gray box represents a word set, with Set 1, 2, and 3 containing 42 words and Set 4 containing 150 words. In BE1, participants were tested on English word Set 1 in Pretest, and learned the same Set 1 words in the Study phase. Set 1, together with Set 2 words were tested in Posttest and Retest. In BE2, an additional Set 3 was included in the Retest phase. BE3 and BE4 were identical to BE1, except that BE3 used Russian words and BE4 used pseudowords. The words in Set 1, 2, and 3 corresponded to “Studied”, “Unstudied”, and “New” words, respectively. EE used the same set of English words throughout the experiment. The selection of words in each word set was randomized across participants. **(B)** Trial procedures of behavioral experiments and auditory stimuli. Each rectangular box along the time axis shows a frame displayed to the participants. Top: Trial procedures for the Pretest, Posttest, and Retest. Each trial started with fixation, followed by simultaneous presentation of the written and auditory word. Then, after seeing the “#” mark, participants indicated the stress position by keypress. The amplitude of the stressed syllable is 20% greater than the other syllables, as indicated by below the spectrogram and schematic RMS amplitude of a word with the stress at the second syllable. Bottom: Study phase procedure. After the display of a written word, the gesture video was presented simultaneously with the audio of the word. **(C)** Videos and conditions in the Study phase. Left: Schematic trajectories of the hand movement in the videos. Right: Frames of the gesture videos. Stressed gestures consisted of three up-and-down movements in front of the body with one movement larger than the other two. The position of the larger movement either matches (*Match* gesture) or mismatches (*Mismatch* gesture) with the stress position in the audio. *Balanced* gesture consisted of three identical movements. Each movement was aligned with each syllable. In the *Static* gesture, the hand remained static throughout the video.

We generated audio of the words using an online text-to-speech tool (Baidu Translate). These audio files were then processed using Adobe Audition CC 2018 software to modify the duration, amplitude, and pitch of the syllables. All syllables were adjusted to 333 ms in duration, resulting in each word of 999 ms. We reduced the pitch variation within each word by adjusting the pitch of each timepoint towards the average pitch while ensuring the audio still sounds natural. We equalized the amplitude of the unstressed syllables to match the average amplitude of the word and increased the amplitude of the stressed syllable to 20% higher than the unstressed syllables (Fig. 1B).

We recorded five videos (Fig. 1C) of manual gestures as visual stimuli during the learning phase. Videos 1 to 4 consisted of three continuous up-and-down manual movements in the vertical plane in front of the torso. In each movement, the hand initially moved downward and then returned upward to the starting position. These videos were synchronized with the audio, temporally aligning each movement with each syllable. Gestures in Videos 1 to 3 were “stressed”, meaning two out of the three movements in each video were standard, while the remaining movement was a larger one with more distance traveling downward. The stressed gestures were further classified into “*Match* gesture” and “*Mismatch* gesture” depending on whether the position of the large movement coincided with the stressed syllable. Video 4 displayed the “*Balanced* gesture”, where all three movements were standard. Video 5 displayed the “*Static* gesture”, where the picture of a static manual gesture in front of the chest was shown.

##### Procedure

Each participant underwent two sessions on two consecutive days. The first session consisted of three phases: Pretest, Study, and Posttest. The second session included a Retest. This experiment utilized a 4 × 2 × 2 mixed design with the gesture type (*Match*, *Mismatch*, *Balanced*, *Static*) being the between-group factor, and the word type (Studied, Unstudied) and test time (Posttest, Retest) being the within-subject factors.

During the Pretest phase, participants were asked to determine the stress position of 42 words randomly selected from the pool of 84 words. There were 14 words for each of the three stress positions. Each trial began with a fixation cross displayed on the center of the screen for 1000 ms, followed by a 999 ms simultaneous presentation of a written word and its auditory presentation. A “#” sign then appeared at the center of the screen, prompting participants to indicate the stress position by keypress. The next trial began immediately after the response without providing any feedback on the response.

During the Study phase, the 42 words that were tested in the Pretest phase (referred to as “Studied words”) were presented again along with gesture videos. Participants engaged in passive listening to the audio and viewing the video during this phase. Each word was studied twice in two consecutive trials. In each trial, after a 1000 ms fixation period and a 1000 ms written word, the audio and a gesture video were simultaneously displayed for 999 ms. Participants were randomly assigned to one of four groups, with 31 participants in each group. Each group was exposed to a different type of gesture video: *Match* group (gesture matched the stress position), *Mismatch* group (gesture mismatched the stress position), *Balanced* group (gesture remained identical), and *Static* group (gesture remained static).

Following a one-minute rest, the Posttest phase began, where participants were tested on both the 42 Studied words and the rest 42 “Unstudied words” from the 84-word pool. The trial procedure for the Posttest phase was identical to that of the Pretest phase.

Approximately 24 hours later, participants returned to the lab to perform the Retest. All 84 words (both Studied and Unstudied) were tested again using the same procedure as the Posttest phase. The presentation order of the words was randomized in all phases and for all participants.

#### 2.2.2 Behavioral Experiment 2

The procedure of BE2 closely resembled BE1, with two exceptions. Firstly, we increased the number of words in Retest from 84 to 126. The additional 42 trisyllabic words used in BE2 were also selected from the COCA word frequency table, with 14 words for each stress position (frequency rank for words that stress on the 1^st^ syllable: *M* = 13396.71, *SD* = 4124.59, *range* = 8436-20199; on the 2^nd^ syllable: *M* = 12987.64 *SD* = 3517.33, *range* = 10001-20165; and on the 3^rd^ syllable: *M* = 12755.14, *SD* = 4056.65, *range* = 8160-19921). The processing methods for the additional audio files remained consistent with BE1. These 42 words were included in Retest in addition to the 84 Studied and Unstudied words as the “New words” that were never presented before the Retest. Secondly, only *Balanced* gesture video was presented during the Study phase.

#### 2.2.3 Behavioral Experiment 3

In BE3, we used Russian words instead of English words. Initially, we compiled a trisyllabic Russian word list from Wiktionary. Then, to ensure that participants were not distracted by unfamiliar speech sounds and word forms, we applied three criteria to shorten the list. We excluded words that: 1) contained letters with no comparable pronunciations in Mandarin or English (Ж, ж, Р, р, Х, х, Ц, ц, Ш, ш, Щ, щ, Ы, ы, Й, й); 2) had large consonant clusters (more than two consecutive consonants); 3) consisted of more than 8 or less than 5 letters. Subsequently, we randomly selected 130 words from the shortened list and then manually selected 84 words by ensuring an equal distribution of 28 words for each stress position.

The audio files of all words were generated by a text-to-speech tool (Google Translate) and then processed by Adobe Audition CC 2022 software to adjust the duration and amplitude of syllables. The duration and amplitude processing methods were consistent with BE1. Moreover, in BE3, we improved pitch control using Praat software. We adjusted the pitch towards the average pitch of each word and limited the pitch variation within a word to less than 60 Hz. Only the videos of *Match* gestures were presented in BE3. The procedure for BE3 was identical to that of the *Match* group in BE1.

#### 2.2.4 Behavioral Experiment 4

In BE4, we substituted the real words with 84 trisyllabic pseudowords generated using VoiceGen (Yang, 2019). Each pseudoword consisted of 3 consonant-vowel syllables. To construct each word, we independently and randomly selected three different consonants from the set /b/, /d/, /p/, /t/, and three different vowels from the set /a/, /e/, /i/, /o/, /u/. The written representation of the words presented to the participants was a direct combination of the letters representing the sounds, such as “datebu”. We then randomly divided the words into three equal groups, with each group assigned one of the three stress positions. The generated audio files were processed using the same method as in BE1. The procedure for BE4 was identical to that of the *Match* group in BE1.

#### 2.2.5 EEG Experiment

In EE, we adapted the paradigm of BE1 for EEG recordings by increasing the number of trials, separating the auditory and written presentation of the word, and introducing randomized intertrial intervals. 150 words selected from the COCA word frequency table were used in this experiment, with 50 words for each stress position (frequency rank for the words that stress on the 1^st^ syllable: *M* = 13674.22, *SD* = 3827.27, *range* = 8436-20199; on the 2^nd^ syllable: *M* = 13367.50, *SD* =3344.26, *range* = 10001-20165; on the 3^rd^ syllable: *M* = 13560.76, *SD* = 4221.09, *range* = 8160-19921). The audio processing methods remained the same as in BE1.

EE consisted of Pretest, Study, and Posttest phases, and only *Match* gestures were presented during the Study phase. In the Pretest, all 150 words were tested. Each trial began with a fixation for a random duration of 400-500 ms, followed by a 1000 ms written presentation of the words. After another random duration of 400-500 ms fixation, the audio was played for 999 ms, while the fixation cross remained on the screen. After a 300 ms blank period, a “#” sign appeared, prompting participants to judge and report the stress position by the keypress. During the Study phase, each trial started with a fixation for a random duration of 400-500 ms. The written word was then displayed for 1000 ms, followed by another random duration of 400-500 ms fixation period. Then, the audio was played simultaneously along with the *Match* gesture video for 999 ms. Each word was studied twice in two consecutive trials, yielding a total of 300 trials during the Study phase. The Posttest followed the same procedure as the Pretest, where all 150 words were tested again. We arranged 3 short breaks throughout the experiment: one between the Pretest phase and Study phase, one after 150 trials (75 words) in the Study phase, and one between the Study phase and the Posttest phase. Each break lasted approximately 2-4 minutes.

### 2.3 Behavioral data acquisition and analysis

The behavioral experiments were carried out in a soundproof room. Participants were instructed to attend to the screen and sound in the headphones. The experiments were controlled using MATLAB 2019a with Psychtoolbox. The visual stimuli were displayed on a Dell 24-inch 2417DG screen with a refresh rate of 144 Hz and a resolution of 2560 × 1440. The audio stimuli were delivered through Sennheiser HD 280 headphones. The keypress responses were recorded using a standard keyboard.

To assess the test performance, we calculated the response accuracy for each participant in each condition. This was done by computing the ratio of correct responses to the total number of responses. As a result, we obtained five accuracies for each participant: Pretest, Unstudied Posttest, Studied Posttest, Unstudied Retest, and Studied Retest. Within each group, we compared the latter four accuracies to the Pretest using two-sided paired t-tests to examine whether the accuracy under each condition differed from the Pretest at a group level.

### 2.4 EEG data acquisition and preprocessing

The EEG experiment was carried out in a soundproof and electromagnetically shielded room. Before the experiment, we verbally familiarized the participants with the general procedure and instructed them to reduce their body, head, and eye movements. The experiment was delivered by the same equipment as used in the behavioral experiments.

EEG signals were recorded from a 32-electrode EEG system (actiChamp system, Brain Products GmbH, Germany). The placement of electrodes on the EasyCap was according to the international 10/20 system. The signals were recorded in single DC mode and sampled at 1000 Hz. Two additional electrodes, horizontal electrooculogram electrodes (HEOG) and vertical electrooculogram electrodes (VEOG), were used to record signals of ocular movements. Electrode impedance was maintained below 10 kΩ. Data were acquired using Brain Vision PyCoder software, online referenced to electrode Cz, and filtered by a 200 Hz low-pass filter and a notch filter at 50 Hz.

Offline processing and analysis of the collected EEG data were conducted using MNE-Python (Gramfort et al., 2014) and EasyEEG (Yang et al., 2018). The data for each participant was band-pass filtered at 0.1-30 Hz. Channels with excessive noise were rejected and interpolated using signals from nearby channels. To eliminate artifacts caused by eye movements, we performed an independent component analysis (ICA) where 25 components were ranked by their correlation with the two electrooculography channels. The artifact components were determined by visual inspection and then manually rejected. The data during the Pretest and Posttest phases were segmented into 1200 ms epochs spanning between 200 ms before and 1000 ms after the auditory stimulus onset. The epochs were baseline corrected by subtracting the average signal during the 200 ms prestimulus period and were re-referenced to the average of all channels. Auto-rejection of epochs where maximum peak-to-peak amplitude exceeded 100 µV, as well as manual rejection of epochs contaminated by eyeblink and movement artifacts, were applied to all epochs.

### 2.5 Temporal domain analysis

Event-related potentials (ERP) were obtained by averaging across epochs for each condition. We computed the global field power (GFP) as the activation index by taking the standard deviation of the ERP responses across all channels at each time point for each participant. The GFP values were averaged across all trials within a condition (6 conditions in total: 3 stress positions in Pretest vs. Posttest). For each individual, the amplitude of the N1 and P2 components was determined by averaging the GFP amplitudes over a 20 ms time window centered around the N1 and P2 peaks, respectively. A paired t-test was then performed to examine whether the amplitude of N1 and P2 differed between the Posttest and Pretest.

### 2.6 Time-frequency analysis

We further conducted a time-frequency analysis to examine the induced rhythmic patterns. We applied the Morlet wavelet transform implemented in the MNE-Python toolbox on the preprocessed epochs to obtain the power and intertrial phase coherence (ITC) of each frequency band at each time point at each channel, with the number of cycles in the wavelet set to frequency/2. ITC was computed based on the following equation (Luo & Poeppel, 2007; Tallon-Baudry et al., 1996):

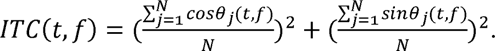

Within each condition, the individual power and ITC averaged for 6 frequency bands: delta (1–2Hz), theta (3-7Hz), alpha (8–12 Hz), low-beta (13–16 Hz), mid-beta (17-20 Hz), and high-beta (21–28 Hz) bands. We quantified the learning-induced changes using a nonparametric spatiotemporal cluster-based permutation test (Maris & Oostenveld, 2007), by comparing the Posttest and Pretest for each stress position, separately for power and ITC. For each stress position, the data combining both Posttest and Pretest trials were randomly partitioned, and a test statistic for each partition was then calculated. This partitioning process was repeated 10000 times to create a null distribution of cluster-level *t* statistics. The *p*-value was determined by the proportion of random partitions whose statistics were larger than the empirical one. The observed clusters were considered significant if *p*-values were below a threshold of 0.05.

## 3 Results

The pretest accuracies in all experiments were above the chance level (0.33) (BE1, *Match*: *M* = 0.54, *SD* = 0.15; BE1, *Mismatch*: *M* = 0.57, *SD* = 0.11; BE1, *Balanced*: *M* = 0.54, *SD* = 0.16; BE1, *Static*: *M* = 0.57, *SD* = 0.14; BE2: *M* = 0.56, *SD* = 0.14; BE3: *M* = 0.41, *SD* = 0.08; EE: *M* = 0.59, *SD* = 0.14; BE4: *M* = 0.43, *SD* = 0.11). These results demonstrated that participants understood the instructions and correctly performed the stress-identification task.

### 3.1 Behavioral Experiment 1

Paired t-tests were carried out to compare the response accuracies in Posttest and Retest with those in Pretest for each group. As shown in Fig. 2, for the group that studied with *Match* gestures, the performance of stress position judgments for the Studied words significantly improved after learning (*Match*, Studied, Posttest: *M* = 0.64, *SD* = 0.17, *t*(30) = 4.26, *p* < 0.001). This improvement lasted until the second day (*Match*, Studied, Retest: *M* = 0.61, *SD* = 0.17, *t*(30) = 2.83, *p* = 0.008). Other types of gestures did not elicit learning effects for the Studied words (*Mismatch*, Studied, Posttest: *M* = 0.53, *SD* = 0.13, *t*(30) = −1.54, *p* = 0.14; *Mismatch*, Studied, Retest: *M* = 0.58, *SD* = 0.14, *t*(30) = 0.45, *p* = 0.66; *Balanced*, Studied, Posttest: *M* = 0.57, *SD* = 0.17, *t*(30) = 1.37, *p* = 0.18; *Balanced*, Studied, Retest: *M* = 0.58, *SD* = 0.17, *t*(30) = 1.96, *p* = 0.06; *Static*, Studied, Posttest: *M* = 0.59, *SD* = 0.17, *t*(30) = 0.81, *p* = 0.42; *Static*, Studied, Retest: *M* = 0.60, *SD* = 0.18, *t*(30) = 1.14, *p* = 0.26). However, the learning effects observed in the *Match* group did not generalize to the Unstudied words (*Match*, Unstudied, Posttest: *M* = 0.55, *SD* = 0.18, *t*(30) = 2.56, *p* = 0.80; *Match*, Unstudied, Retest: *M* = 0.56, *SD* = 0.18, *t*(30) = 0.87, *p* = 0.39). These results suggested that gestures help in learning lexical stress when the features between gestures and auditory stimuli are congruent.

**Figure 2.**
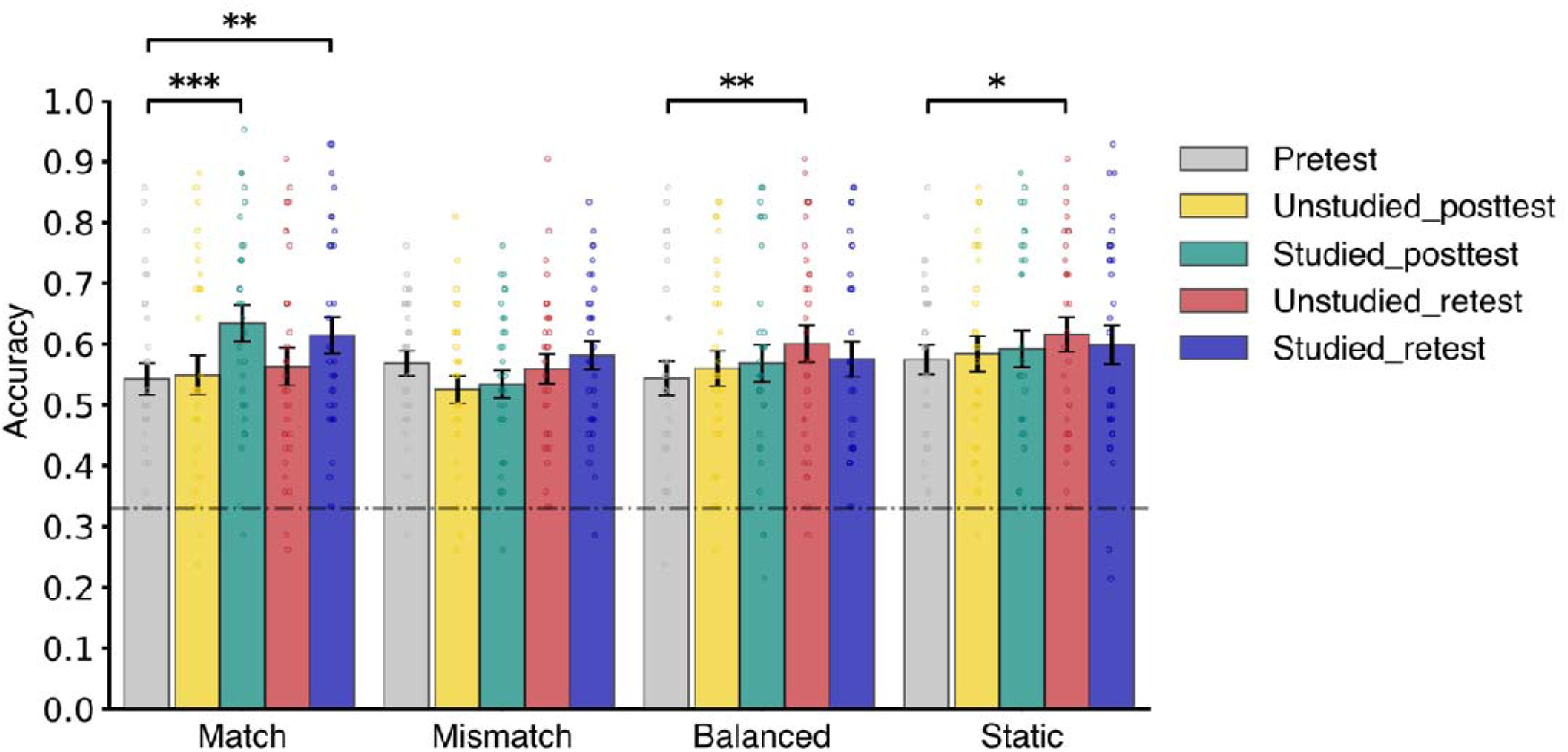
Results of BE1. Mean response accuracies with individual data of the 4 groups (*Match*, *Mismatch*, *Balanced*, and *Static*) in BE1. Each bar represents the mean accuracy within a group. Each small circle is an individual accuracy. In each group, the five bars from left to right respectively represent the accuracy for the conditions of Pretest, Unstudied words in Posttest, Studied words in Posttest, Unstudied words in Retest, and Studied words in Retest. Compared to the Pretest, the performance of stress position judgment is significantly better in the Posttest and the Retest of Studied words for the *Match* group, as well as in the Retest of Unstudied words for both *Balanced* and *Static* groups. The horizontal dashed line indicates the chance performance (0.33). The error bars indicate ± SEM. Asterisks indicate the significance level of the accuracy differences evaluated by paired t-tests, ∗*p* < 0.05, ∗∗*p* < 0.01, ∗∗∗*p* < 0.001.

Following our initial hypothesis, we did not expect any changes after learning in the *Balanced* group and *Static* group, because these two types of gestures conveyed no information about the stress position. Surprisingly, the *Balanced* and *Static* groups showed increased accuracy in Retest compared to Pretest for Unstudied words (*Balanced*, Unstudied, Retest: *M* = 0.60, *SD* = 0.18, *t*(30) = 2.85, *p* = 0.008; *Static*, Unstudied, Retest: *M* = 0.62, *SD* = 0.16, *t*(30) = 2.13, *p* = 0.04). No other conditions showed significant changes (*Mismatch*, Unstudied, Posttest: *M* = 0.53, *SD* = 0.13, *t*(30) = −1.85, *p* = 0.07; *Mismatch*, Unstudied, Retest: *M* = 0.56, *SD* = 0.13, *t*(30) = −0.36, *p* = 0.72; *Balanced*, Unstudied, Posttest: *M* = 0.56, *SD* = 0.16, *t*(30) = 0.92, *p* = 0.36; *Static*, Unstudied, Posttest: *M* = 0.58, *SD* = 0.17, *t*(30) = 0.51, *p* = 0.62). These results hinted at possible generalization elicited by *Balanced* gestures, but the observed effects for the Unstudied words in Retest could also be due to the exposure of the same set of words during the Posttest. We therefore further carried out BE2 to understand these unexpected results.

### 3.2 Behavioral Experiment 2

In BE1, the *Balanced* group showed improved response accuracy for Unstudied words in the Retest, suggesting a potential generalization of learning. However, since all words in Retest were previously presented in Posttest, it is possible that the observed effects were due to the repeated exposure of stimuli (test-retest effects) rather than true generalization. Generalization should yield performance improvements even for new words that are not previously encountered. To test whether the *Balanced* gesture induced true generalization, we repeated the *Balanced* group experiment on another group of participants, with a set of New words added to the Retest.

We successfully replicated the increased accuracy for the Unstudied words in Retest observed in BE1 (Fig. 3; *M* = 0.62, *SD* = 0.16, *t*(31) = 3.16 *p* = 0.003). Crucially, the response accuracy for the New words also significantly increased after learning (*M* = 0.62, *SD* = 0.17, *t*(31) = 3.48, *p* = 0.002). These results indicated that *Balanced* gestures elicited generalization. Additionally, for this group of participants, the performance also improved across the rest conditions (Posttest, Unstudied: *M* = 0.60, *SD* = 0.14, *t*(31) = 2.32, *p* = 0.027; Posttest, Studied: *M* = 0.59, *SD* = 0.14, *t*(31) = 2.45, *p* = 0.019; Retest, Studied: *M* = 0.62, *SD* = 0.17, *t*(31) = 3.70, *p* < 0.001).

**Figure 3.**
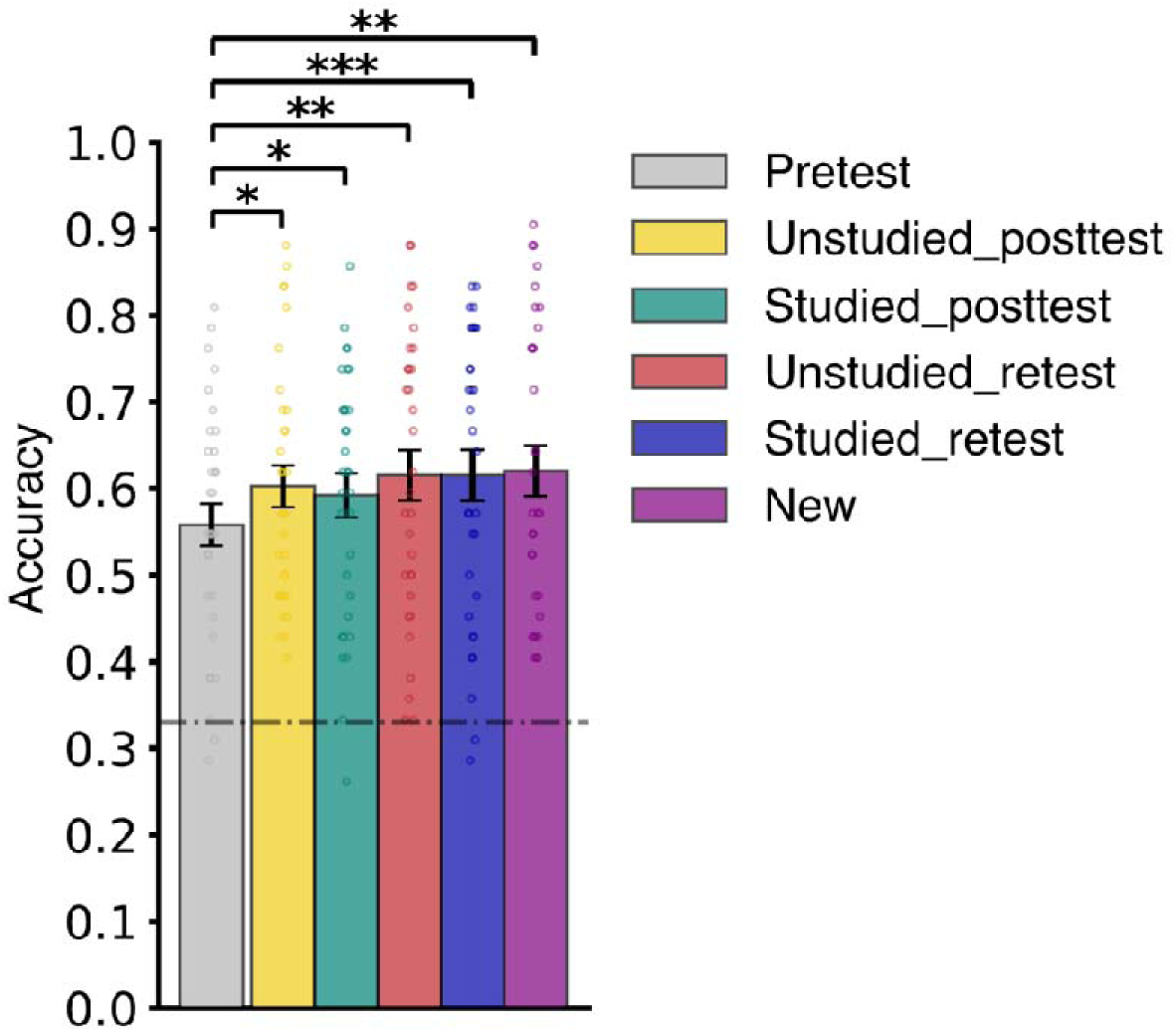
Results of BE2. Mean response accuracies with individual data in BE2 (*Balanced* gesture). The plotting conventions are identical to Fig. 2. After learning with *Balanced* gestures, the response accuracies improved for all conditions, including the New words in the Retest that had never appeared in previous phases.

### 3.3 Behavioral Experiment 3

The goal of BE3 was to investigate whether lexical information is necessary for *Match* gestures to facilitate the learning of lexical stress. Because all participants reported above-moderate English proficiency, the word stimuli in BE1 provided lexical information to the participants, including semantic, morphological, and phonological/orthographic information. To remove lexical information, we substituted the English words with Russian words in BE3. Post-experiment interviews confirmed that almost all participants had zero previous exposure to Russian. Only one participant had previous experience of Russian, but his test performance was similar to other participants. Therefore, we could assume that participants had no previous lexical knowledge about the Russian word stimuli.

Replicating the learning effects for English words, *Match* gestures significantly improved accuracy for Studied Russian words on the Posttest (Fig. 4; *M* = 0.46, *SD* = 0.09, *t*(30) = 2.50, *p* = 0.018) and Retest (*M* = 0.45, *SD* = 0.09, *t*(30) = 2.39, *p* = 0.023). Interestingly, contrasting with English stimuli, the learning effect of Russian was generalized to the Unstudied words in Posttest (*M* = 0.46, *SD* = 0.09, *t*(30) = 2.67, *p* = 0.012). However, the generalization effect did not last in Retest (*M* = 0.43, *SD* = 0.08, *t*(30) = 0.97, *p* = 0.34). These results demonstrated that *Match* gestures could facilitate lexical stress learning without lexical information, although the generalization may be temporary.

**Figure 4.**
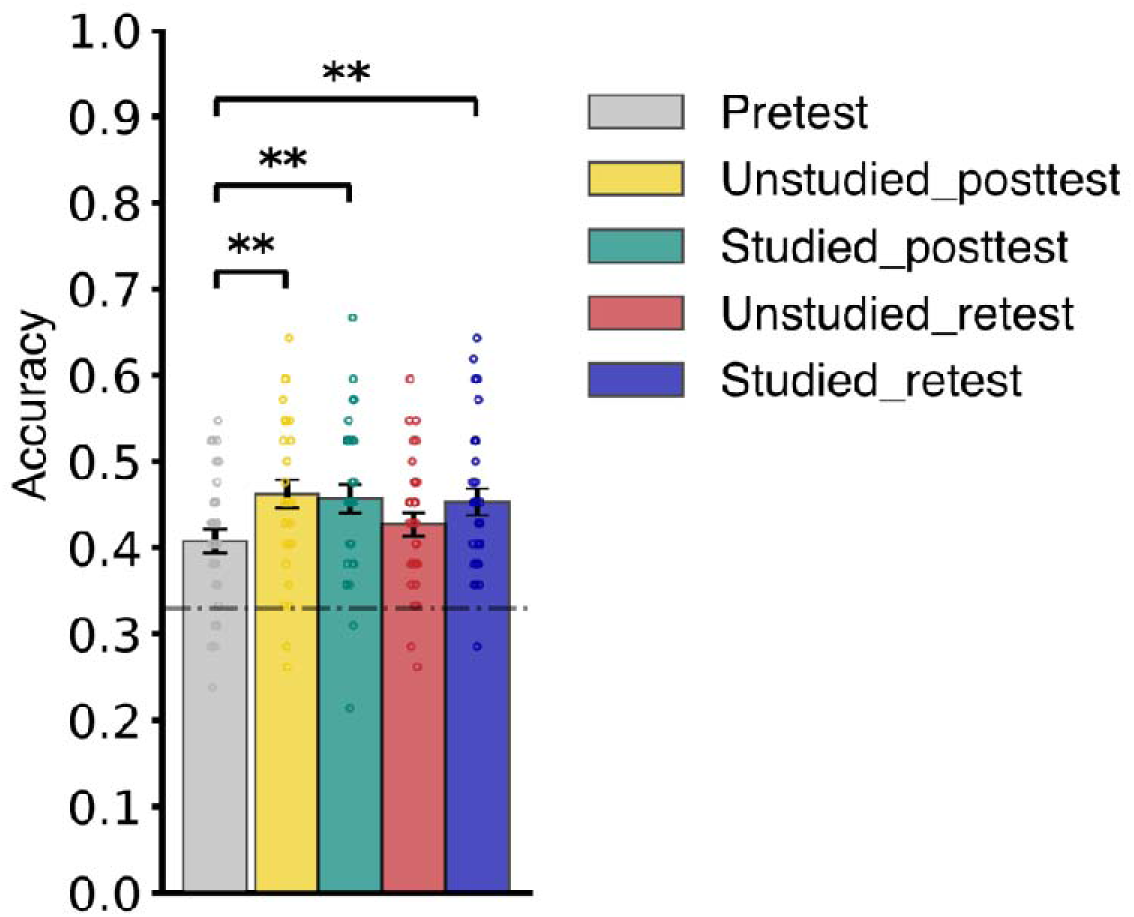
Results of BE3. Mean response accuracies with individual data in BE3 (Russian words). The plotting conventions are identical to Fig. 2. The performance improved for both Unstudied and Studied words in Posttest, but the improvement lasted in Retest only for the Studied words but not the Unstudied words.

### 3.4 Behavioral Experiment 4

Are the observed learning effects simply due to the sensitivity increase to acoustic features (e.g. amplitude) rather than learning of phonological features of stress? We investigated this question in BE4. In BE4, we performed the *Match* group experiment again, replacing the real words with randomly generated pseudowords, which did not follow the phonotactic regularities of real words.

No significant change in test performance was found (Fig. 5; Unstudied, Posttest: *M* = 0.45, *SD* = 0.11, *t*(30) = 0.64, *p* = 0.53; Studied, Retest: *M* = 0.42, *SD* = 0.12, *t*(30) = −0.70, *p* = 0.49; Unstudied, Retest: *M* = 0.43, *SD* = 0.12, *t*(30) = −0.03, *p* = 0.97; Studied, Retest: *M* = 0.44, *SD* = 0.11, *t*(30) = 0.42, *p* = 0.68). These results indicated that the *Match* gestures could not improve learning the pseudoword stress given the same studying and testing procedures as in BE1 and BE3, suggesting the observed learning effects cannot simply explained by the improvement of sensitivity to acoustic features.

**Figure 5.**
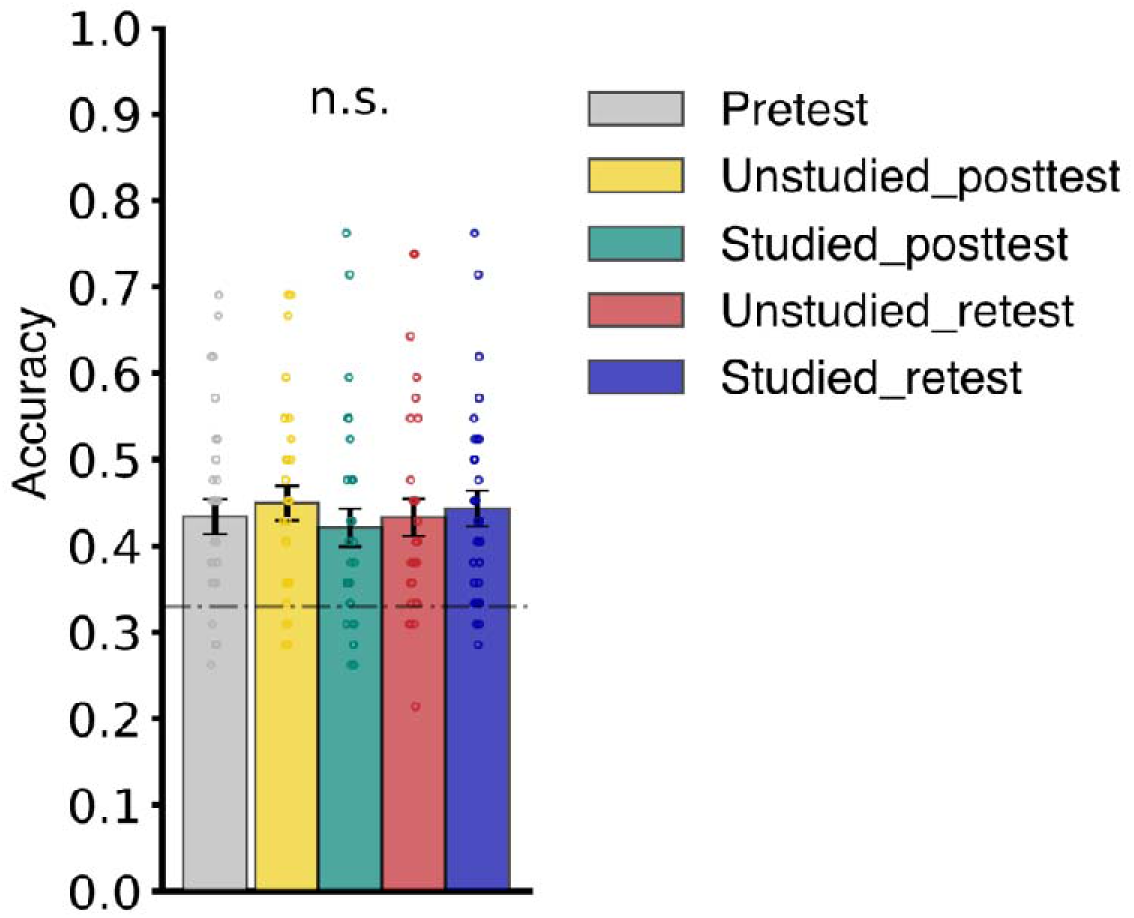
Results of BE4. Mean response accuracies with individual data in BE4 (pseudowords). The plotting conventions are identical to Fig. 2. No significant changes in response accuracy were found.

### 3.5 EEG Experiment

#### Behavioral results

The EEG experiment replicated the behavioral results of the Studied words of the BE1 *Match* group. Paired *t*-test revealed that the response accuracy in Posttest (*M* = 0.68, *SD* = 0.13) significantly increased after learning (*t*(20) = 5.10, *p* < 0.001), indicating that the *Match* gesture also facilitated learning in the EEG experiment.

#### EEG results

Because we have observed the learning effect at the behavioral level, our next step was to investigate neural correlates of this effect by comparing the EEG responses in Posttest with those in Pretest. For all stress positions, we observed typical auditory responses, N1 and P2 components with central frontal and parietal topographic distribution, respectively (Fig. 6). To statistically determine whether the N1 and P2 amplitudes changed after learning, we computed peak amplitude values of GFP at around 100 ms and 200 ms for each subject, representing their individual N1 and P2 amplitudes. We carried out paired t-tests to examine the group-level changes in these amplitudes before and after learning. The results showed that the P2 amplitude in response to the first syllable-stressed words significantly decreased in the Posttest (*M* = 2.72, *SD* = 0.73) compared to the Pretest (*M* = 3.14, *SD* = 1.11; *t*(20) = −2.44, *p* = 0.024). However, the P2 amplitude did not show significant changes for the second syllable-stressed words (Pretest: *M* = 2.96, *SD* = 1.31; Posttest: *M* = 2.55, *SD* = 0.86; *t*(20) = −1.92, *p* = 0.070) or the third syllable-stressed words (Pretest: *M* = 3.12, *SD* = 1.00; Posttest: *M* = 2.74, *SD* = 0.91; *t*(20) = −2.01, *p* = 0.058). No significant changes in N1 amplitude were found for words with stress on the first syllable (Pretest: *M* = 2.26, *SD* = 0.60; Posttest: *M* = 2.20, *SD* = 0.62; *t*(20) = −0.41, *p* = 0.68), the second syllable (Pretest: *M* = 2.23, *SD* = 0.65; Posttest: *M* = 2.20, *SD* = 0.55; *t*(20) =-1.17, *p* = 0.26), or the third syllable (Pretest: *M* = 2.31, *SD* = 0.68; Posttest: *M* = 2.25, *SD* = 0.70; *t*(20) = −0.38, *p* = 0.71).

**Figure 6.**
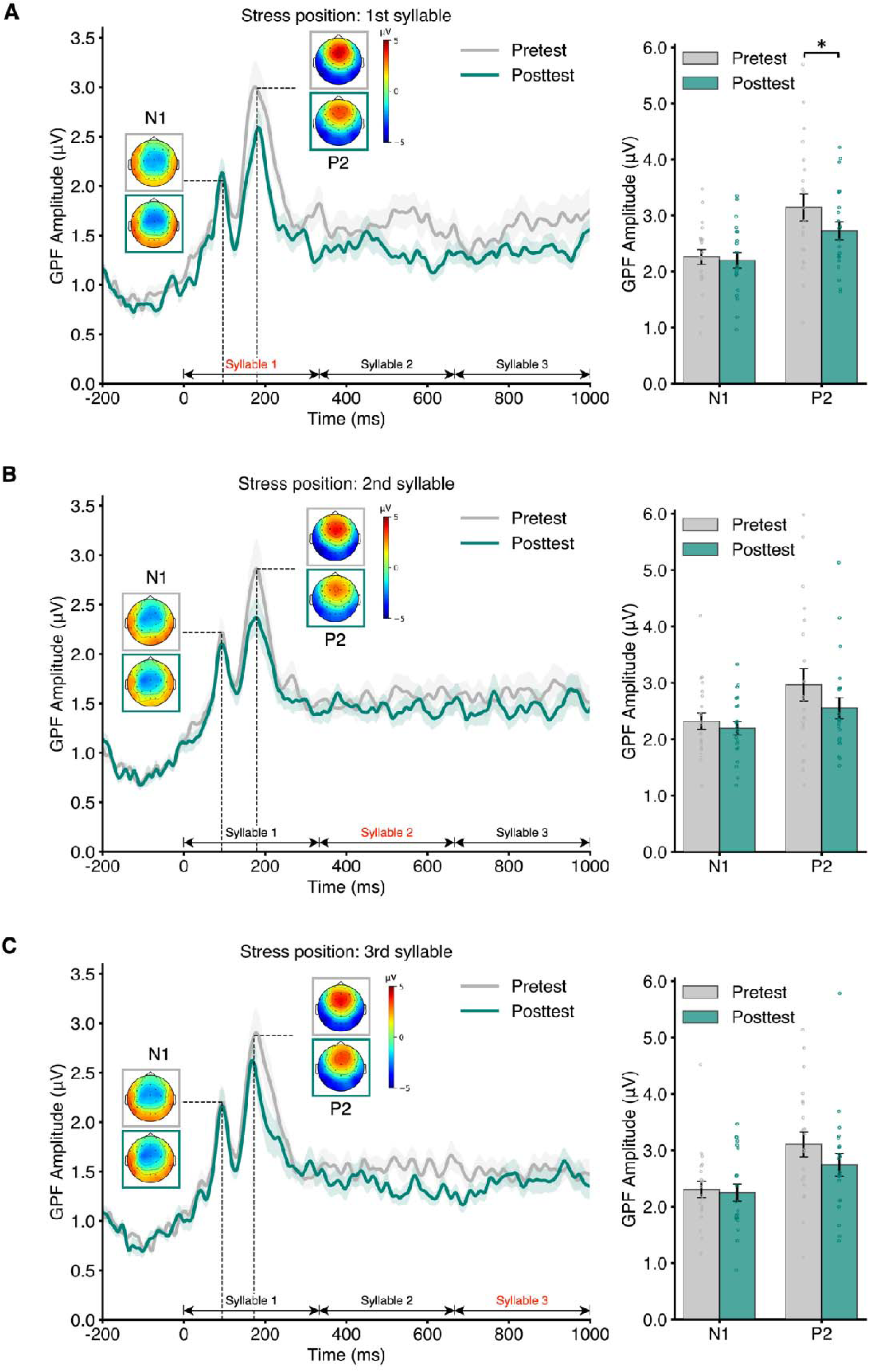
Results of GFP waveforms and N1 and P2 components. Left: GFP evoked by the auditory stimuli averaged across all participants. The grey line and the green line represent Pretest and Posttest GFPs respectively. The shades around the lines represented ± SEM. Time zero corresponds to the auditory stimulus onset time, and the time windows of the syllables are marked above the time axis. The vertical dashed lines indicate the peaking latencies of N1 and P2 components averaged between Pretest and Posttest, pointing to the N1 and P2 topographies in Pretest (grey box) and Posttest (green box). Right: Mean N1 and P2 amplitudes with individual data in Pretest and Posttest. The error bars indicate ± SEM. The asterisk indicates the significance level of the amplitude difference in Pretest and Posttest evaluated by paired t-test. ∗*p* < 0.05, ∗∗*p* < 0.01, ∗∗∗*p* < 0.001. The three panels respectively plot responses to words with stress on **(A)** the first syllable, **(B)** the second syllable, and **(C)** the third syllable. The P2 amplitude showed a trend to decrease after learning for all three groups of words, but the difference reached significance for only the words with stress on the first syllable.

These temporal domain results are mostly consistent with our hypothesis. The absence of effects in the early N1 component suggests that the learning effects are not primarily driven by the sensitivity changes to the acoustic features, consistent with the null results observed in BE4. Moreover, the P2 effects reached significance only for the first syllable-stressed words but not for the other words with later stress positions. One reason is that the temporal analysis is time-locked to the stimulus onset that induces prominent peaks of early perceptual ERP components. The learning effects of the stress at the first position temporally align and reflect in the P2 component. Whereas, the learning effects of later stress positions may not be apparent in the relatively weaker ERP responses at later latencies, making it difficult to directly detect the effects elicited by the latter two syllables using temporal domain analysis. While the behavioral improvement of stress perception was observed across words with all stress positions in behavioral experiments, the learning effects should occur at each stress position. Therefore, we performed a time-frequency analysis that can be carried out without time-locking to the onset of stimuli to uncover neural responses to stress learning at various positions by examining the induced rhythmic patterns.

For all three stress positions, spatiotemporal cluster-based permutation tests revealed significant enhancement of power (first: *p* = 0.018; second: *p* = 0.0209; third: *p* = 0.0496) and suppression of ITC (first: *p* = 0.0165; second: *p* = 0.0003; third: *p* = 0.0469) in the theta frequency band (3-7 Hz) after learning (Fig. 7). Importantly, the latencies of the significant clusters varied as a function of the stress positions. For power, the significant cluster appeared early for words with stress on the first syllable (−200 to 959 ms), later for words with stress on the second syllable (219 to 1000 ms), and even later for words with stress on the third syllable (373 to 964 ms). A similar trend was observed in ITC: the latencies for the significant clusters shifted to later time points as the stress positions moved from the first syllable (−131 to 290 ms) to the second syllable (−98 to 562 ms) and to the third syllable (366 to 695 ms). That the dynamics of learning effects corresponded to the stress positions suggested that gestures facilitate stress learning in a position-specific manner.

**Figure 7.**
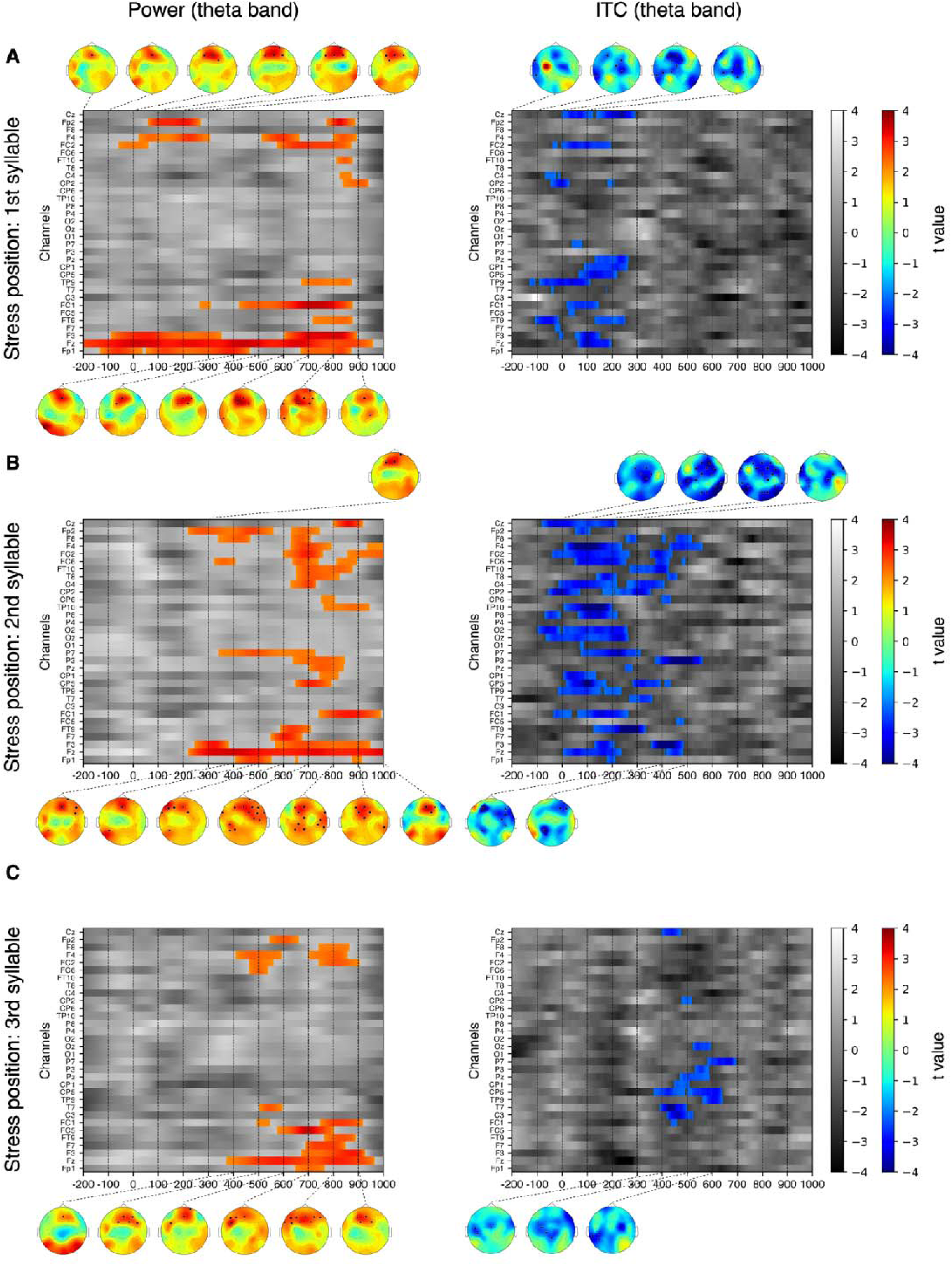
Results of time-frequency analysis in the theta frequency band (3-7 Hz). Left: Theta band power. Right: Theta band ITC. Each image shows the *t* value at each channel and each time point of comparing the power or ITC in Posttest to Pretest by spatiotemporal cluster-based permutation tests (Posttest minus Pretest). The *t* value is depicted using a colored scale if it belongs to a significant cluster, and grayscale if it does not. Topographies of *t* values are plotted alongside using the consistent colored scale every 100 ms where significant clusters were observed, with significant channels marked with black squares. The three panels respectively plot responses to words with stress on **(A)** the first syllable, **(B)** the second syllable, and **(C)** the third syllable.

Moreover, the significant clusters displayed distinct topographic patterns. The significant channels that showed power differences were primarily located in frontal regions with a few extending to the parietal regions (Fig. 7). Whereas, those for ITC differences were more widely distributed with a focus on parietal regions. These topographies indicated that gestures may induce plasticity over the auditory cortices via the influence of the frontal-parietal network.

## 4 Discussion

The present study investigated the effects of manual gestures on learning lexical stress. In four behavioral and one EEG experiments, consistent evidence supports that gestures facilitate the learning of lexical stress by modulating perceptual neural responses to corresponding stress positions. The gesture facilitation effects were observed in both languages that are familiar (English) and unknown (Russian) to participants but absent in pseudowords that lack phonotactic regularities, suggesting that the gestures facilitate the learning at the level of suprasegmental phonological features. Moreover, gestures that match the amplitude variations in the lexical stress facilitate the learning of the studied words, whereas gestures that only match the segment timing but not in amplitude generalize the learning effects to unstudied stimuli. These results suggest that the gesture facilitation effects depend on the specificity of cross-modal feature mapping at the levels of target abstract mental constructs.

Although the connection between gestures and language acquisition is widely recognized, previous studies have rarely tested lexical-phonological speech attributes. Specifically, no studies have successfully shown that gestures could affect the learning of lexical stress (Chang et al., 2014; Jesse & Mitterer, 2011). Bridging this gap, our study presents evidence demonstrating the benefit of gestures in lexical stress learning. In our learning paradigm, participants learned the lexical stress of foreign languages by observing the up-and-down manual movements synchronized with audio recordings of spoken words. We have demonstrated that gesture-accompanied learning significantly enhances the stress perception of the studied words in both English and Russian (BE1-3, Figs. 2-4).

One of the core factors that may ground the emergence of such a cross-modal facilitation effect is the congruent relation between gesture and auditory stimuli. In our study, we established this congruency by aligning the spatial trajectory of gestures with the acoustic speech envelope, which represents the amplitude profile of lexical stress. This relation is both ecologically valid and aligns with participants’ prior understanding of lexical stress as a strengthened syllable. Consequently, participants might find it relatively effortless to map the gestural features onto the auditory space, resulting in significant learning effects despite a low training load. Previous studies that failed to demonstrate an effect of gestures on lexical stress perception may partially be due to the ineffective establishment of the mapping between gestural and acoustic features. For example, in the Jesse and Mitterer (2011) study, a drawing of a disembodied hand was used to depict pointing gestures, which may have appeared unnatural and thus difficult to integrate with the auditory stimuli.

Results in BE1 (Fig. 2) support the hypothesized function of congruency in gesture-accompanied learning. Moreover, the results demonstrate possible constraints of cross-modal congruency on generalization. The four types of gestures formed a spectrum of congruency, ranging from more to less congruent: *Match* gestures (specifically congruent) > *Balanced* gestures (generally congruent, because merely the segment timing was aligned) > *Static* gestures (generally incongruent, could also be understood as neutral or irrelevant) > *Mismatch* gestures (specifically incongruent). Interestingly, we indeed observed a corresponding spectrum of learning effects. *Match* gestures, at the one end of the spectrum, profoundly enhanced the learning of the studied words, whereas *Mismatch* gestures, at the other end, had no significant effects. A deviation from our initial hypothesis was that *Balanced* gestures, situated in the middle of the congruency spectrum, also facilitated learning. This observation aligns with a recent report on beat gestures (similar to our *Balanced* gesture) influencing the perception of lexical stress (Bosker & Peeters, 2021).

Importantly, we observed a distinction in the effect patterns between generally congruent gestures and specifically congruent gestures – only *Balanced* (generally congruent) but not *Match* (specifically congruent) gestures elicited generalization. The generalization effects of *Balanced* gestures were replicated in BE2 after ruling out the test-retest factor (Fig. 3). This spectrum of the effect patterns observed across variations of the relationship between the visual and auditory stimuli highlights the importance of the nature (generality vs specificity) of congruency in the across-modal mapping for learning.

Moreover, we found generalization when learning Russian words with *Match* gestures (Fig. 4), similar to the generalization when learning English words with *Balanced* gestures (Figs. 2-3). These results suggest that the amount of information trained may constrain generalization in learning. Learning involves reducing outcome uncertainty by distinguishing informative cues from uninformative cues (Rescorla, 1988). Consistent with this theory, a recent study on language learning found that the complexity of cues influences generalization (Vujović et al., 2021). Specifically, the presence of complex information before simple information enhances item learning, while the presence of simple information before complex information promotes generalization (Vujović et al., 2021). In the present study, specifically congruent (*Match*) gesture provides more complex information, explicitly indicating the stress position. Consequently, participants tend to focus on and memorize the stress position of specific words, resulting in a stronger item-learning effect, but no generalization effect. On the other hand, generally congruent (*Balanced*) gestures provide simple information about segment timing but not stress position, facilitating generalization. Meanwhile, generalization was also observed when we reduced the complexity of the auditory stimuli. For the participants in this study, Russian stimuli are simpler than English stimuli because of the absence of lexical information. As a result, fewer features of the Russian stimuli could be extracted and utilized for item memorization. This shift in learning focus may enhance the ability to recognize stress patterns within the speech stream, leading to the identification of stress in both studied and unstudied words. That is, training on too much information may incline overfitting – multiple aspects of information are good for memorization but could reduce generalization.

Our findings in behavioral experiments (Fig. 2-5) collaboratively indicate that the facilitation effects of gestures on lexical stress learning primarily occur at the phonological level. According to the levels of processing theory (Craik & Lockhart, 1972), information encoded at a deeper level is easier to remember. Therefore, it is not surprising that gestures can aid in learning words that involve high-level knowledge, as participants can rely on their existing lexical knowledge to encode the stress position of each token. The intriguing part of this study is that by using Russian stimuli, we have demonstrated that gesture can still benefit learning at the phonological level for suprasegmental features without the aid of any lexical information. Moreover, we did not observe the effects in BE4 where the stimuli were simple concatenations of consonant-vowel syllables with no phonotactic regularities, which made judgments of stress positions only based on the differences of the acoustic amplitude of each syllable segment. This suggests that the learning effects of gestures are not a general enhancement of auditory sensitivity.

To our knowledge, our study provides the first neural evidence of the impact of gestures on learning suprasegmental speech attributes. For the ERP responses, we observed the decreased response magnitude for the learned items at the perceptual component of P2 (Fig. 6). These results support the hypothesis that the learning effects occur at the phonological level. Moreover, the effects were absent in N1, which is consistent with behavioral results that the learning effects were not due to increased sensitivity to lower-level acoustic attributes. The ERP results only revealed the learning effects for the items with stress at the beginning but not later positions. This is presumably because the ERP responses are more sensitive to the onset of stimuli and hence have more power to reveal the effects at the first position.

More importantly, the complementary spectral analyses demonstrate that gesture-accompanied learning specifically modulated the auditory neural responses at the corresponding stress positions We observed an increase of theta band (3 – 7 Hz) power after gesture-accompanied learning, consistent with a previous study showing that theta power elicited by words that appear with gestures is greater compared with those appear without gestures (Biau et al., 2015). Note that the onset of the modulation effect preceded the actual onset of the stressed syllable, which is consistent with previous studies showing that the preceding activation potentially is associated with the change in the expectation of auditory stimuli (Arnal & Giraud, 2012; Biau et al., 2015). Crucially, the modulation effect was localized to the latency of the stress position, rather than spanning the entire stimulus duration (Fig. 7). Such temporal alignment between the modulation effect and stress position was also found in ITC, further supporting that the mechanism of lexical stress learning is selective feature imprinting rather than whole stimulus storage (Goldstone, 1998). These neural results suggest that the visual system (visual motion of biological movement) can directly or indirectly affect the auditory system in the early stages of perception, thereby facilitating the learning of lexical stress.

Notably, we report a decrease, instead of an increase, in theta band ITC after learning, despite the improvement of stress perception performance. This finding is opposite to previous auditory perceptual learning studies where the ITC of low-frequency oscillations increases along with improved behavioral performance (Andrillon et al., 2015; Luo et al., 2013; Ringer et al., 2023). This might indicate a distinct mechanism between learning lexical stress and basic acoustic patterns. One possible explanation is that lexical stress is a suprasegmental feature that spans across time. The brain may sample across multiple time points to detect the stress pattern. Learning may expand the sampling range and diversify the sampling point to increase the chance and efficiency of detecting the patterns of lexical stress, thereby leading to less phase coherent responses across trials.

In summary, our study provides consistent evidence that gestures facilitate the learning of lexical stress by modulating auditory neural responses. The facilitation effects occur at the phonological level and the generalization depends on the cross-modal specificity of feature mapping. Our findings highlight the functional role of gestures in enhancing speech learning, suggesting practical implications for language teaching and learning.

## Acknowledgments

We thank Junchun Yang for her assistance in data collection. This study was supported by the National Natural Science Foundation of China 32071099 and 32271101, Program of Introducing Talents of Discipline to Universities, Base B16018, and NYU Shanghai Boost Fund.

## Author contributions

D. Yang and X. Tian conceived the study. T. Zhan, D. Yang, and X. Tian designed the experiments. T. Zhan, D. Yang, and R. Wu performed the experiments and analyzed the data. T. Zhan and X. Tian wrote the paper. X. Tian supervised the study.

